# HMGB1 assists in overcoming cisplatin resistance in chemoresistant human ovarian cancer cells

**DOI:** 10.1101/2025.09.25.678487

**Authors:** Van Huynh, Guliang Wang, Anirban Mukherjee, Karen Vasquez

## Abstract

Cisplatin is one of the most effective chemotherapeutic agents used in the treatment of ovarian cancer. However, the frequent development of cisplatin resistance remains a significant limitation, leading to therapeutic failure and poor patient outcomes. Cisplatin cytotoxicity is attributed to the generation of toxic DNA lesions, which can be recognized and processed by a variety of proteins, including the high mobility group box 1 (HMGB1) protein. HMGB1 is a multifunctional protein, which is involved in chromatin remodeling and multiple DNA damage repair pathways. In this study, we investigated the role of HMGB1 in modulating cisplatin sensitivity in human ovarian cancer cells. Using cisplatin-sensitive and cisplatin-resistant human ovarian cancer cell lines, we employed siRNA-mediated HMGB1 knockdown to assess its impact on the cellular responses to cisplatin treatment. In clonogenic survival assays, HMGB1 depletion resulted in a significant reduction in colony formation in cisplatin-resistant cells upon cisplatin exposure, compared with non-targeting siRNA treated cells. Additionally, HMGB1 inhibition significantly enhanced cisplatin-induced apoptosis in the cisplatin-resistant cells. Mechanistically, HMGB1-depleted cells exhibited altered DNA damage responses via modulation of ATM/CHK2 and ATR/CHK1 activity following cisplatin treatment. Notably, DNA immunoblot and modified alkaline comet assay results demonstrated that HMGB1 depletion stimulated cisplatin-DNA adduct formation and impaired the removal of cisplatin-DNA adducts, particularly in the cisplatin-resistant cells. Collectively, these findings uncover novel functions of HMGB1 in mediating cisplatin sensitivity, emphasizing its potential as a therapeutic target to overcome cisplatin resistance in ovarian cancer.

## Introduction

Ovarian cancer is the ninth most commonly diagnosed cancer among women and remains the deadliest gynecologic malignancy in the United States, with an estimated 20,890 new cases and approximately 12,730 deaths in 2025 [1]. Most ovarian cancer patients are diagnosed at advanced stages, resulting in a low five-year survival rate of ∼31% for distant-stage disease [1]. Chemotherapy, particularly platinum-based regimens, remains one of the standard treatments for ovarian cancer [2, 3]. Cisplatin [cis-diamminedichloroplatinum (II)] is used effectively to treat several human tumor types, including ovarian tumors [4–6]. Although cisplatin is initially effective, more than 70% of advanced-stage ovarian cancer patients experience relapse within two years after treatment [7]. Most of these relapsed cases develop resistance to cisplatin, which is a significant factor contributing to treatment failure and poor long-term outcomes [3, 7–9]. Therefore, understanding the molecular mechanisms of cisplatin resistance is crucial for identifying potential therapeutic targets and strategies to improve treatment outcomes.

Cisplatin can interact with purine bases generating DNA intrastrand and interstrand crosslinks (ICLs) [10, 11]. These covalently linked ICLs are highly toxic lesions, preventing strand separation during DNA transcription, replication, and repair, and have been attributed to the majority of cisplatin cytotoxicity [12, 13]. Multiple mechanisms have been implicated in cisplatin resistance, including dysfunctions in drug transport, drug inactivation, increased DNA damage responses and repair activities, upregulation of anti-apoptotic activity, and/or tumor microenvironment alterations [14, 15]. Since cisplatin can target the DNA, the ability of cancer cells to respond to and remove cisplatin-induced DNA lesions is an important underlying mechanism of resistance [16]. The cellular responses to cisplatin-induced DNA damage are complex, initiated by the activation of critical kinases in the DNA damage response (DDR) network, including ataxia telangiectasia mutated (ATM) and ATM and Rad-3-related (ATR), CHK1 and CHK2 through phosphorylation [17–20]. DDR activation, in turn, triggers multiple downstream cellular events, such as cell cycle arrest, DNA repair to maintain genomic integrity or apoptosis if damage is extensive [17, 19]. While cisplatin intrastrand lesions are mainly removed by nucleotide excision repair (NER) [21, 22], cisplatin ICL removal involves multiple repair proteins/pathways, including Fanconi Anemia (FA), NER, mismatch repair (MMR), translesion synthesis (TLS), and homologous recombination (HR) [23, 24]. Thus, modulating the efficiency of the DDR and DNA damage repair activities has been targeted to resensitize chemoresistant cancer cells to cisplatin [24, 25].

The high mobility group box 1 (HMGB1) protein, a member of the HMGB protein family, is a non-histone architectural nuclear protein highly expressed across human cell types [26, 27]. Structurally, HMGB1 contains two DNA binding domains, the A-box and B-box, and the C-terminal acidic tail. Boxes A and B promote DNA binding and bending, and the C-acidic tail regulates the binding affinity [28–30]. The biological functions of HMGB1 are versatile depending on its cellular location [26, 31]. Intracellular HMGB1 has a high binding affinity to distorted DNA structures such as triplex DNA, cruciform DNA, bent DNA, and chemotherapeutic-induced DNA damage such as cisplatin DNA adducts [29, 32, 33]. Studies have shown the potential diagnostic and prognostic roles of HMGB1 in multiple cancers, including ovarian cancer since its overexpression has been indicated and is often associated with poor treatment outcomes due to chemoresistance [34–36]. Whether HMGB1 impacts cisplatin sensitivity in chemoresistant ovarian cancer via modulating the processing of cisplatin-modified DNA lesion removal remains unclear. However, HMGB1 has been reported to play a role in multiple DNA repair pathways related to the repair of cisplatin DNA adducts, such as NER, MMR, and HR [26, 37]. Utilizing triplex technology to direct a site-specific psoralen ICL in DNA substrates via psoralen-conjugated triplex-forming oligonucleotides (TFOs), we showed that HMGB1 binds ICLs specifically and with high affinity and promotes ICL recognition and processing via cooperating with NER proteins, such as XPC-RAD23B and RPA, and XPA [38–41]. Furthermore, it has been shown that HMGB1 preferentially binds to 1,2-d(GpG) and d(ApG) cisplatin intrastrand DNA substrates [29, 42]. Confocal microscopy imaging directly revealed HMGB1-Pt-DNA complexes in cells [43], and acetylated or phosphorylated forms of HMGB1, together with other HMGB family members, were found to recognize and bind to cisplatin-damaged DNA [44]. These findings implicate HMGB1 in the processing of cisplatin-induced DNA adducts.

Importantly, we recently demonstrated that inhibiting HMGB3, another member of the HMGB family that shares a high level of similarity to HMGB1, enhances cisplatin sensitivity in chemoresistant human ovarian cancer cells by influencing apoptosis and ATR/CHK1 in the DDR [45]. HMGB3 and HMGB1 share high similarities in sequence and structure but appear to have distinct biological functions [46–48]. Thus, in this study, we aim to explore the impact of HMGB1 on cisplatin sensitivity in chemoresistant human ovarian cancer cells and the underlying mechanisms involved. Using cisplatin-sensitive (A2780) and cisplatin-resistant (CP70) human ovarian cancer cell lines, we show that HMGB1 depletion followed by cisplatin treatment in CP70 cells resulted in a decrease in colony formation as well as an increase in the apoptotic cell population. Furthermore, we found that DDR activity was altered following cisplatin treatment as a function of HMGB1 depletion. In addition, depletion of HMGB1 facilitated cisplatin DNA adduct formation and diminished the efficiency of the removal of cisplatin-DNA adducts.

## Materials and Methods

### Cell lines and culture conditions

Human ovarian cancer cell lines, including cisplatin-sensitive A2780 (RRID:CVCL_0134) and cisplatin-resistant A2780/CP70 (CP70, RRID:CVCL_0135), were purchased from the American Type Culture Collection (Manassas, VA, USA) with short tandem repeat profiling (STR) verification for authentication. Cell lines were grown in RPMI-1640 medium supplemented with 10% fetal bovine serum (FBS) in a 37°C humidified incubator with 5% CO_2_. Confluent cells (80% - 90%) were dissociated by 0.25% trypsin EDTA and split at 1:5 - 1:3 (A2780) or 1:4 - 1:3 (CP70) dilutions for sub-culturing. CP70 cells were grown in cisplatin-containing media at a final concentration of 1 µM every three passages to maintain drug resistance. Experiments were performed when cells reached 70% - 80% density for both cell lines at the 3^rd^ passage.

### Cisplatin treatment, MTT assay, and IC50 determination

A 3.3 mM cisplatin stock solution was freshly made for each experiment, as previously described [45]. Briefly, 1 mg of cisplatin was added into 1 mL of PBS containing 140 mM NaCl solution, followed by vortexing at high speed at room temperature for 5 minutes and incubating at 37°C for 20 minutes. The stock can be stored at 4°C, protected from light for up to one month. Cell viability was evaluated by using the MTT (3[4,5-dimethylthiazol-2-yl]-2,5-diphenyltetrazoliumbromide) assay using the CellTiter 96™ Nonradioactive Cell Proliferation Assay kit (Promega) according to the manufacturer’s instructions. Briefly, A2780 and CP70 cells were seeded in triplicate in a 96-well plate at a density of 15,000 cells/well and incubated at 37°C for attachment and growth. After 24 hours, cells were treated with 200 µL cell culture medium containing cisplatin at half-serially diluted concentrations from 100 μM for 72 hrs at 37°C. Cells incubated with a drug-free culture medium and the medium-only wells were included as controls and blanks, respectively. Following cisplatin treatment, the media was removed, and the cells were washed with warm PBS twice. After adding 100 µL fresh cell culture media and 15 µL premixed MTT dye to the wells, the cells were incubated at 37°C for 3 hrs in the dark. Postincubation, solubilization solution was added to the culture wells at a 1:1 volume ratio, and the plate was gently shaken overnight at room temperature in the dark. The absorbance at 570 nm was then recorded using a 96-well plate reader (Synergy H1; BioTek Instruments, Inc.). The cisplatin concentrations that inhibit 50% of the cell viability were determined as the IC50 for each cell line.

### siRNA treatment

HMGB1 expression was transiently inhibited using siRNA as previously described [40] with slight modifications. A siGENOME SMARTpool containing four different siRNA sequences against HMGB1 (Cat. No. M-018981-01-0020) and siGENOME non-targeting siRNA#2 (Cat. No. D-001210-02-20) were purchased from Dharmacon and resuspended in 1X siRNA buffer made from 5X siRNA buffer (Dharmacon). Reverse and forward transfections were performed for 500,000 cells (A2780) and 850,000 cells (CP70) using Lipofectamine RNAiMAX (Invitrogen) according to the manufacturer’s protocol. The final concentrations of siRNAs and RNAiMAX were optimized for each cell line, including A2780 (50 nM and 9 µL) and CP70 (30 nM and 7 µL). During reverse transfection, siRNA and RNAiMAX were diluted in separate tubes containing 250 µL Opti-MEM Reduced Serum medium (Life Technologies). After adding the siRNA solution in a dropwise manner into the RNAiMAX solution, the mixture was incubated at room temperature for 5 - 10 minutes (extending over 10 minutes will decrease the efficiency). The prepared cell suspensions at the indicated cell numbers in antibiotic-free culture media were added to the reagent mixture. After adding Optimem to the final volume of 2.5 mL, the cells were plated in 60 mm dishes and incubated at 37°C, 5% CO_2_. Then, the media was changed after 24 hrs, and the cells were allowed to recover for another 24 hrs. During the forward transfection, the reagent mixtures were prepared similarly and filled to a final volume of 2.5 mL with Optimem. After removing the media, the cells were exposed to the transfection reagent mixtures for 24 hrs. Following medium replacement, the cells were cultured for 24 hrs before being collected to check knockdown efficiency or for subsequent experiments.

### Immunoblotting

Protein expression levels were determined by western blot analysis as previously described [49] with modifications. At different time points during the experiments, the cells were harvested using cell scrapers, followed by centrifugation at 13000 rpm at 4°C for 10 minutes. The cell pellets were stored at -80°C before subjecting to SDS-PAGE and western blot analysis. Briefly, the cell pellets were mixed and incubated with cold RIPA buffer (50 mM Tris-HCl, pH 7.4, 150 mM NaCl, 0.1% SDS, 1.0% NP-40, 1% Na-deoxycholate) supplemented with protease inhibitor cocktail (cOmplete^™^, Roche) and phosphatase inhibitor cocktail (PhosSTOP^™^, Roche) for 1 hr. The lysis mixtures were then sonicated for 20 seconds, followed by 20-second intervals on ice, repeated 6 times using a water bath sonicator. The cell lysates were stored at -20°C or used directly for the downstream steps. The protein concentration in cell lysates was identified using the Pierce™ BCA Protein Assay kit (23225, Thermo, USA) and diluted to the working concentration (45 µg) using lysis buffer before mixing 1:1 by volume with 2X Laemmli sample buffer (Bio-Rad, CAT#1610737) and denaturing at 95°C for 10 minutes. After cooling to room temperature, the extracted protein samples were loaded onto 4–15% Mini-PROTEAN® TGX™ Precast Protein Gels (Bio-Rad, CAT#4561086), followed by the gel electrophoresis in 1X Tris-Glycin-SDS buffer at 60V for 15 minutes and 200V for 30 minutes. The proteins were then transferred to 0.2 µm nitrocellulose membranes (Bio-Rad, CAT#1704270) by using the Trans-Blot Turbo Transfer System (Bio-Rad, CAT#1704150) at 2.5 A for 15 minutes. The membranes were then washed with 1X PBST (Phosphate Buffered Saline with 0.1% Tween 20) for 5 minutes three times, followed by a blocking step with 3.7% non-fat dried milk (NFDM - Bio-Rad, CAT#1706404XTU) in PBST for at least 1 hr at room temperature and overnight incubation with primary antibodies at 4°C against Phospho-ATM, Phospho-ATR, Phospho-CHK2, Phospho-CHK1, ATM, ATR, CHK1, CHK2, HMGB1, and β-Actin as the internal loading control (**Supplementary Table 1**). The membranes were then washed with 1X PBST for 5 minutes three times and incubated with Horseradish Peroxidase (HRP)-conjugated secondary antibodies against rabbit and mouse at 1:5000 dilution for at least 1 hr at room temperature. All the incubations and washes were performed with a rotator. Proteins were developed with Clarity Western ECL Substrate (Bio-Rad, CAT#1705061) and imaged using a ChemiDoc XRS+ Imager (Bio-Rad). The band densitometric intensities were analyzed using Image J software (NIH, version 2.14.0/1.54f, RRID:SCR_003070).

### Clonogenic survival assay

Clonogenic survival assays were performed as described [45] following siRNA treatments with HMGB1-targeted siRNA (siHMGB1) and non-targeting siRNA (siNT) in cisplatin-resistant CP70 cells, as indicated previously. Briefly, A2780, CP70, and siRNA-transfected CP70 cells were incubated with culture media containing cisplatin at a concentration 4 times lower than the IC50 identified for CP70. Non-treated A2780 and CP70 were included as controls. After 72 hrs, the cells were counted using an automated cell counter (Invitrogen, CAT#A49866), replated at the density of 1000 cells in 6 mL media in 60 mm dishes, and cultured at 37°C, 5% CO_2_ for 10-14 days for colony formation. Then the media was removed, the colonies were washed, fixed with 95% ethanol for 10 minutes at room temperature, and allowed to dry overnight. For colony visualization, the cells were stained with 0.125% crystal violet solution for 30 minutes at room temperature. Colonies of ≥50 cells were manually counted. Data were shown as the absolute number of colony counts and represented as the mean ± SEM value from four independent experiments. Original two-way ANOVA with Sidak correction was performed for statistical significance (p-value <0.05 *, <0.01 **, <0.001 ***, <0.0001 ****).

### Cell cycle analysis

A2780 and CP70 cells were transfected with siRNAs according to the protocol described above. Non-treated A2780 and CP70 were used as controls. For drug treatment, the cells were incubated with the desired concentration of cisplatin at 37°C in 5% CO_2_ for 72 hrs. Following cisplatin treatment, the cells were trypsinized, counted, centrifuged at 1000 rpm at 4°C for 5 minutes, and washed with cold PBS two times. For fixation, the cells were resuspended in 200 µL - 500 µL cold PBS, then added dropwise into tubes containing 5 mL of chilled 70% ethanol while gently vortexing, and stored at -20°C overnight. The fixed cells were pelleted at 1500 rpm at 4°C for 10 minutes, and washed with cold PBS. Prior to the fluorescence-activated cell sorting by flow cytometry, the cells were resuspended in a staining solution (20 μg/mL Propidium Iodide, 0.5% Triton-X, and 20 μg/mL RNase A in PBS). The single-cell mixtures were obtained by filtering through the snap cap with a 35 µm nylon mesh into tubes and then incubated for 1 hr at 37°C, followed by flow cytometric analysis using the BD Accuri C6 system (RRID:SCR_01959). Cell cycle profiles were generated to determine the sub-G1 cell populations (%) using the FlowJo software (RRID:SCR_008520). Experiments were performed in at least three independent replicates.

### Slot blot assay

Wildtype and siRNA-treated A2780 and CP70 cells were exposed to cisplatin media at the desired treatment concentration for 24 hrs. Following cisplatin treatment, the cells were washed with warm PBS and incubated with fresh media to allow time for DNA repair. The examined samples in each group included T0, T24, and T96 (collected at 0, 24, and 96 hrs following cisplatin treatment, respectively) and non-treated controls. At the indicated time points, the cells were washed, scraped, pelleted, and stored at -20°C. During the processing step, the cell pellets were thawed and resuspended in a solution containing 200 µL PBS, 4 µL RNase A (100 mg/mL) (Qiagen, CAT#19101), and 20 µL Proteinase K (Qiagen). The slot blot procedure was performed as described (DOI: 10.21769/BioProtoc.1453) [50] with modifications. Briefly, genomic DNA was isolated using the DNeasy Blood & Tissue Kit (Qiagen, CAT#69506) according to the manufacturer’s protocol. DNA concentrations were determined using a Nanodrop (ThermoFisher) and diluted with distilled water. Samples containing 600 ng of genomic DNA were denatured at 100° C for 10 minutes and immediately transferred to ice, followed by adding an equal volume of cold 2 M ammonium acetate (pH 7.0) to stabilize the single-stranded DNA. The denatured DNA samples were then spotted onto a cellulose membrane using the Bio-Dot SF Microfiltration apparatus system (Bio-Rad) according to the manufacturer’s protocol. The membrane was baked at 80°C for 2 hours in a hybridization oven to allow DNA fixation. After rehydration in 1X PBST, the membrane was blocked in 3.7% NFDM in 1X PBST for at least 1 hour at room temperature. Following the blocking, the membrane was probed with anti-cisplatin modified DNA primary antibody (Abcam CAT# ab103261, RRID:AB_10715243)) at a 1:15000 dilution in 3.7% NFDR in PBST overnight at 4°C. The membrane was then washed three times in PBST, followed by incubation with anti-rat HRP secondary antibody (Cell Signaling Technology, CAT#7077, RRID:AB_10694715) at a 1:5000 dilution in 3.7% NFDM in PBST for at least 2 hours at room temperature. After washing three times with PBST, the membrane was incubated with the Clarity Western ECL Chemiluminescent reagent (Bio-Rad, CAT#1705061) for 10 minutes. The signals were detected and visualized using the Bio-Rad GelDoc imaging system. The membranes were then briefly washed with PBST and stained with Sybergold at a 1:10000 dilution in PBST for 1 hour at room temperature and destained in PBST overnight, followed by signal detection for a DNA loading control. The band intensities were quantified using Image J software (NIH, version 2.14.0/1.54f). Statistical analyses were performed using GraphPad Prism software (RRID:SCR_002798). Data were represented as the mean ± SEM value from at least three independent experiments. Original two-way ANOVA with Sidak correction was performed for statistical significance (p-value <0.05 *, <0.01 **, <0.001 ***, <0.0001 ****).

### Alkaline comet assay

The cisplatin concentration and the treatment time course were empirically determined in wild-type ovarian cancer cells. The cells in 60 mm dishes were incubated with 2.5 mL of 100 µM cisplatin media for 1 hour. Following cisplatin treatment, cells were washed with warm PBS twice and recovered in fresh media at 37°C to allow time for DNA repair. The samples examined in each group included T10, T24, T48, and T72 (collected at 10, 24, 48, and 72 hours post-treatment, respectively) and non-treated controls. T10 samples were determined as the time point with the highest induced cisplatin ICL levels. T24, T48, and T72 were included to evaluate the timing of repair. CP70 cells in the siNT and siHMGB1-treated groups underwent cisplatin treatment and were collected according to the above protocol. The sample labeling in each group included H_2_O_2_-cis (cisplatin-treated samples with H_2_O_2_ treatment), non-cis (non-treated control samples), H_2_O_2_-non-cis (non-treated control samples with H_2_O_2_ treatment), and cis (cisplatin-treated samples). During collection, the cell numbers were obtained for each sample, and the cells were frozen at a 10^6^ cells/mL density in media containing 10% (vol/vol) DMSO and 10% (vol/vol) FBS at -80 °C in a Mr. Frosty box until the downstream processing step. The alkaline comet procedure was developed based on the described protocol (DOI: 10.1007/978-1-0716-0323-9_7) [51] with modifications. Briefly, the frosted end of the microscope slides were precoated with 1% agarose (Sigma-Aldrich, CAT#A0169) in distilled water and dried overnight at room temperature. A 0.8% low melting point agarose (Sigma-Aldrich, CAT#A9414) in 1X PBS mixture was prepared and cooled in a 37°C water bath. For each sample, two gels were prepared in two separate slides. One slide was exposed to 20 µM hydrogen peroxide (H_2_O_2_) for 15 minutes to induce DNA breaks, and the other was not treated. Each gel contained a cell density of 2 x 10^5^ cells. After thawing in a 37°C water bath, the calculated cell volumes were transferred to a new centrifuge tube, followed by a washing step with 1 mL cold PBS at 9,000 rpm in a cold centrifuge for 5 minutes. The supernatants were discarded, and the cell pellets were placed on ice. 2-3 pre-coated slides were labeled and placed on a metal tray on an ice box. After gently mixing the cell pellets with 0.8% low melting point agarose solution, the agarose-cell mixtures were placed in a 37°C heater during the gel casting to prevent gel solidification. Then 60 µL of the mixtures were transferred onto the precoated slides to make one gel. The 20×20 mm coverslips were placed on the top, and the gels were allowed to solidify on ice for 10 minutes. Once the gels were formed, the coverslips were removed. The process continued the same way for the rest of the samples. The gel-embedded cells on designated slides were placed in a rack and submerged in a box containing 20 µM H_2_O_2_ solution for 15 minutes, followed by a brief wash in 1XPBS. The immediate subsequent lysis step was performed by incubating in the cold lysis buffer (2.5 M NaCl, 0.1 M Na_2_EDTA, 10 mM Tris base, pH 10, and freshly added 1% Triton X-100 and 10% DMSO) overnight at 4°C. After the lysis step, the slides were briefly washed in 1X cold PBS before transferring to a flat-bed electrophoresis tank with the frosted ends toward the anode and submerged in the ice-cold alkali buffer (0.3 M NaOH and 1 mM Na_2_EDTA, pH 12.5) for 30 minutes at 4°C in the dark to allow DNA unwinding. Following alkaline treatment, the electrophoreses were conducted at 1 V/cm for 25 minutes at 4°C in the dark. After electrophoreses, the slides were neutralized in 1X cold PSB for 10 minutes, followed by a washing step with distilled water for 10 minutes. The slides were then dehydrated by incubating with 70% Ethanol (EtOH) for 10 minutes and 100% EtOH for 15 minutes and dried overnight at room temperature in the dark. During the staining step, the slides were rehydrated in water for 30 minutes before incubating with Syber gold (1:10000) in 1X TE solution for 30 minutes. The slides were then washed with water for 20 minutes and allowed to air dry overnight in the dark. Next, 20 µL of water was added, followed by a coverslip placement to each gel immediately before the comet visualization using a Zeiss AxioObserver Z1 inverted fluorescence microscope at x10 magnification. The images were acquired by Zen microscopy software (Zeiss, RRID:SCR_013672). At least 100 comets were scored per sample. The comet analyses were performed using the CometScore 2.0 software. The Olive tail moment (TM) parameter was calculated by multiplying the percentage of total tail DNA by the distance between the intensity centers of the head and the tail regions, as previously described [52]. The TM of the H_2_O_2_-non-cis sample and the H_2_O_2_-cis samples at each time point were normalized to the TM of the non-treated control samples. ICLs in the H_2_O_2_-cis samples inhibited the mobility of the H_2_O_2-_induced fragmented DNA, causing a decrease in TM (DTM) compared to the H_2_O_2_-non-cis samples. The levels of ICLs in the H_2_O_2_-cis samples at every time point were directly proportional to the percentage DTM at that time point compared to the H_2_O_2_-non-cis samples [53] and were calculated using the following formula:

% DTM = [1 – (TM_H2O2-cis_ – TM_non-cis_)/(TM_H2O2-non-cis_ – TM_non-cis_)] × 100

ICL repair was evaluated by identifying the percentages of ICL unhooking at 10, 24, 48, and 72 hours (T10, T24, T48, and T72) post-cisplatin incubation in a drug-free medium, in which the ICL level at T10 was the highest, and the DTM at T10 was set at 100%. The ICL unhooking percentages at T24, as an example, were calculated by the formula:

% ICL unhooking at T24 = [(% DTM at T10 - % DTM at T24) / % DTM at T10] x 100

### Statistical analysis

GraphPad Prism software (version 9) was used for statistical analyses and data visualization. In all graphs, the data were represented as means ± SEM (standard error of the means). All experiments were performed in at least three independent replicates. The student’s t-test or original two-way ANOVA with Sidak’s post-hoc test for multiple comparisons correction was performed to determine statistically significance differences between groups. A p-value <0.05 was considered statistically significant. Detailed statistical information is indicated in the figure legends.

## Results

### HMGB1 depletion enhances cisplatin sensitivity in chemoresistant human ovarian cancer cells

Human cisplatin-sensitive A2780 and cisplatin-resistant CP70 ovarian cancer cell lines were used for this study. The cell lines were tested for cisplatin sensitivity using an MTT assay, and the IC50 of the cisplatin-sensitive CP70 cells was found to be ∼ tenfold higher than that of the cisplatin-sensitive A2780 cells. To investigate the impact of HMGB1 depletion on cisplatin sensitivity in cisplatin-resistant human ovarian cancer cells, we performed clonogenic assays (**Figure 1A**). CP70 cells were treated with siRNA specific to HMGB1 before being exposed to cisplatin for 72 hours. Colony formation analysis was carried out 10-14 days post cisplatin treatment. Western blot analysis indicated >90% reduction in HMGB1 expression resulting from two siRNA treatments (**Figure 1B**). The cisplatin concentration used was ∼4 times lower than the IC50 value determined for CP70 cells. We found that the A2780 cells were sensitive to cisplatin, which resulted in the absence of colony formation. There was a minor reduction in the number of colonies in the CP70 cells and the control siNT-treated CP70 cells. Remarkably, we observed that the colony number after cisplatin treatment was significantly decreased in the HMGB1-depleted CP70 cells (21%) compared to the control (56%) (**Figure 1C, D**). These results demonstrate that HMGB1 depletion impaired clonogenic growth and enhanced the cisplatin sensitivity in chemoresistant cells.

**Figure 1.**
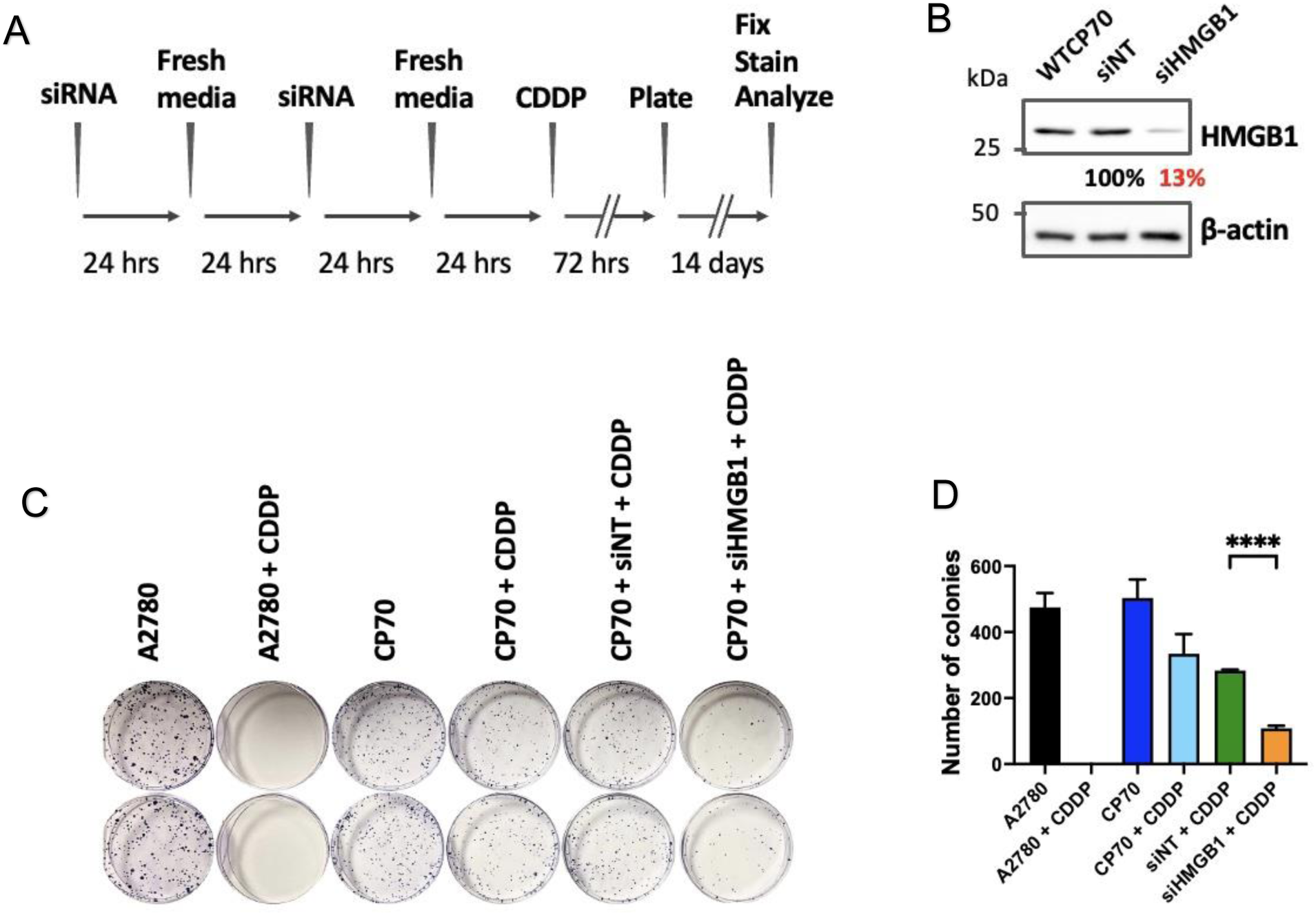
HMGB1 depletion enhances cisplatin sensitivity in chemoresistant human ovarian cancer cells. **A**, Schematic outline of the experimental design for the clonogenic assays. HMGB1 expression in cisplatin-resistant cells (CP70) was transiently silenced using an siRNA (50 nM) directed against HMGB1. Control cells were transfected with a non-targeting siRNA (50 nM). Following incubation with 1.5 µM cisplatin (∼ 3-fold lower than the IC50) for 72 hours, 1000 cells were plated in 60 mm dishes and allowed to grow at 37°C, in a 5% CO_2_ incubator. After 2 weeks, the effect of HMGB1 depletion on colony formation was analyzed. CDDP, cisplatin. **B**, Western blotting was performed to determine the HMGB1 protein level for knockdown efficiency. More than 85% of HMGB1 was depleted as an average from three independent experiments. Samples were collected at the time point of plating. WTCP70, wild-type CP70; siNT, non-targeting siRNA; siHMGB1, siRNA against HMGB1; β-actin, loading control protein. **C**, Representative image of a clonogenic survival assay in each group. Cells were fixed with 100% EtOH for 10 minutes, stained with 0.125% crystal violet in dH20 for 20 minutes, and colonies of ≥50 cells were counted. A2780, cisplatin-sensitive human ovarian cancer cells. **D**, Counted colonies of the treatment groups were quantified and represented in the bar graphs. Four independent experiments were performed. Data are mean ± SEM and were analyzed by Student’s t-test, **p-value <0.01, n = 4).

### Cisplatin-induced apoptosis was significantly increased in HMGB1-depleted ovarian cancer cells

Apoptosis is a critical cellular response for cisplatin cytotoxicity; thus, we evaluated the effects of HMGB1 depletion on cisplatin-induced apoptosis in human ovarian cancer cells. We treated the A2780 and CP70 cells with siRNA to deplete HMGB1, and then treated the cells with cisplatin (∼4 times lower than the IC50 identified for CP70 cells) for 72 hours. Flow cytometry with PI staining was utilized to analyze the sub-G1 apoptotic cell population across the samples (**Figure 2A**). HMGB1 levels were reduced by ∼85% and 90% in the A2780 and CP70 cells, respectively, as assessed by western blotting (**Figure 2B**). A representative histogram of cell cycle analysis by FlowJo software for these two cell lines showed a substantial sub-G1 population in **Figure 2C, D** (average of ∼40% from >3 repeats) in the control A2780 cells following cisplatin treatment, indicating cisplatin-induced apoptosis. A significant increase to 45.2% in apoptotic cells (**Figure 2C, D,** average of ∼49% from >3 repeats) was observed in the HMGB1-depleted A2780 cells. In the chemoresistant CP70 cells, cisplatin treatment resulted in ∼10% of the cells in the sub-G1 population in the siNT-treated sample. Notably, we found that in the HMGB1-depleted CP70 cells, the sub-G1 population was significantly increased to ∼40% in response to cisplatin treatment (**Figure 2E, F**). This finding indicates that HMGB1 depletion facilitated cisplatin-induced apoptosis in both the A2780 and CP70 cells, consistent with the clonogenic survival data. Together, these results suggest a role for HMGB1 in resensitizing chemoresistant ovarian cancer cells to cisplatin.

**Figure 2.**
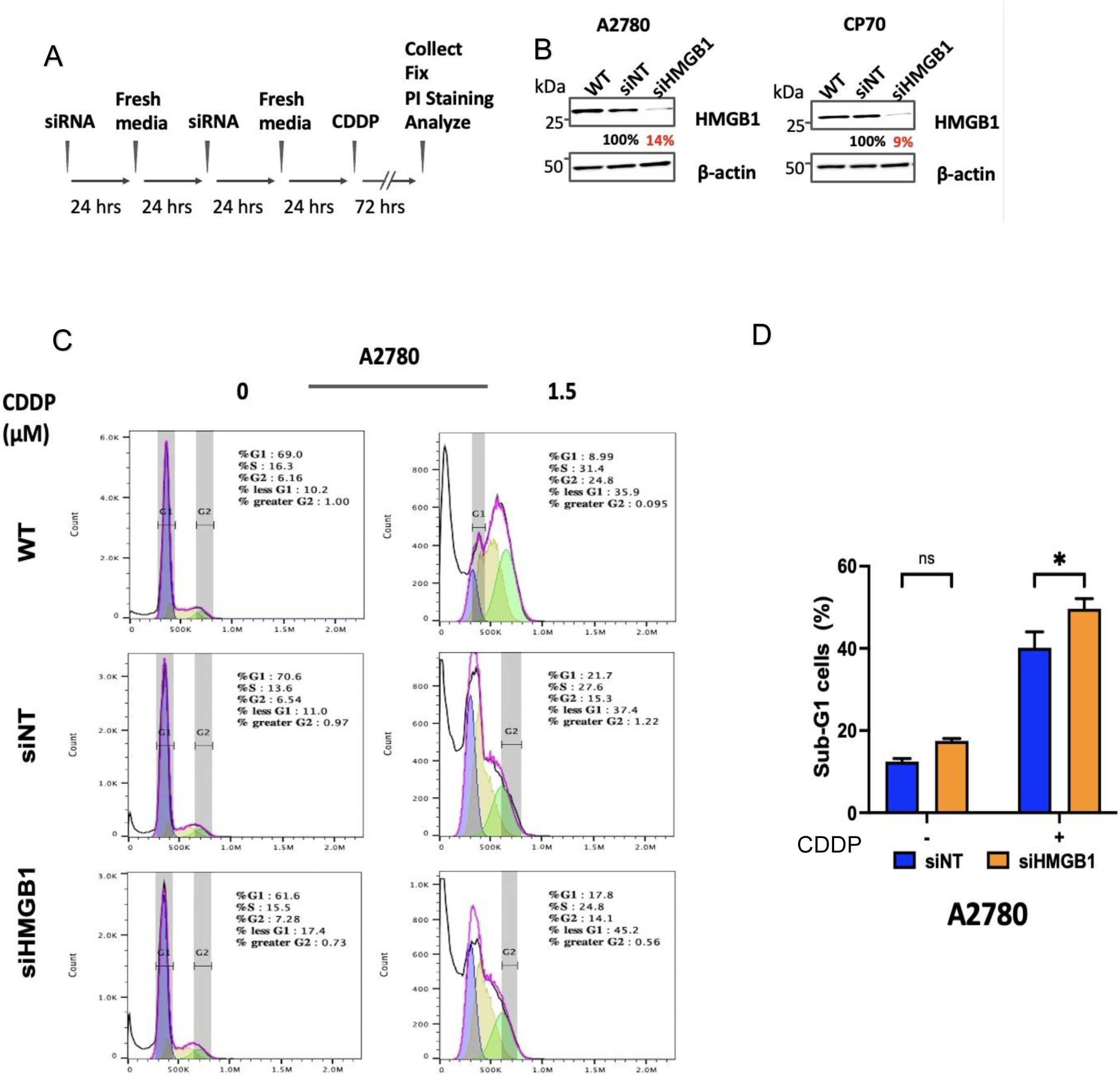
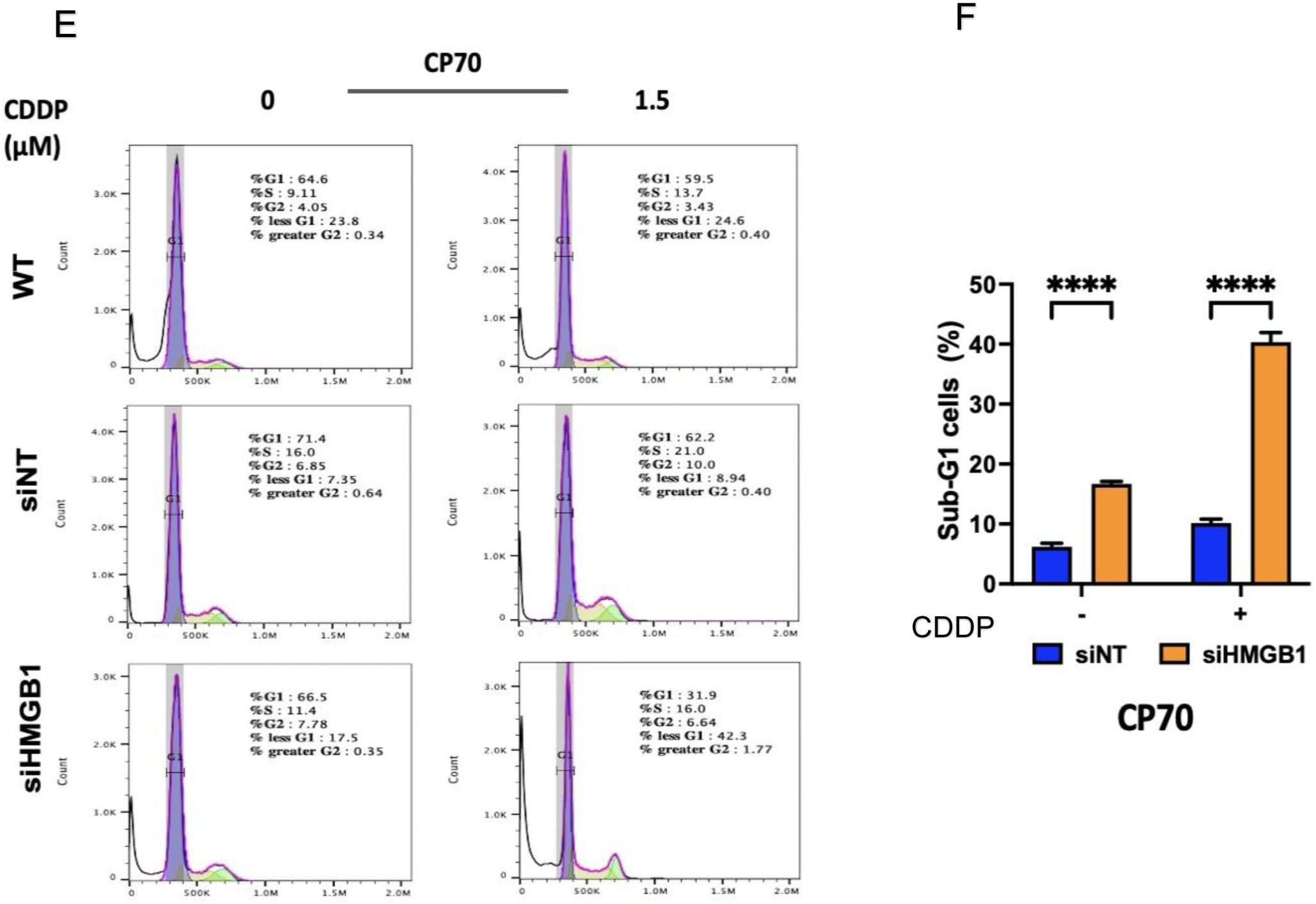
Cell cycle analysis demonstrates an increase in apoptosis following cisplatin treatment as a function of HMGB1 depletion in human ovarian cancer cells. **A**, The experimental design for the cell cycle analysis is illustrated in the schematic. A2780 and CP70 cells were transfected with either non-targeting siRNA or an HMGB1 siRNA (50 nM). Following exposure to 1.5 µM cisplatin (CDDP) for 72 hours, the cells were collected, counted, and fixed with 75% EtOH. Fixed cells were washed and subjected to propidium iodide (PI) staining for 2 hours. Flow cytometry was performed to detect apoptotic cells via the sub-G1 population. **B**, HMGB1 depletion was evaluated using western blotting. β-actin served as the loading control. The average HMGB1 knockdown efficiency from three independent experiments was 85% and 90% for A2780 and CP70, respectively after 2 siRNA treatments. A2780, cisplatin-sensitive A2780; CP70, cisplatin-resistant A2780CP70; siNT, non-targeting siRNA; siHMGB1, siRNA against HMGB1. **C**, **E**, Representative flow cytometric histograms depict the percentage cells across different cell cycle phases for A2780 and CP70, respectively. The apoptotic cell population is defined as sub-G1 cells. **D**, **F**, HMGB1 inhibition significantly enhanced apoptosis in response to 1.5 μM cisplatin treatment in A2780 and CP70 cells, respectively. Quantification of the sub-G1 cell population in each sample was analyzed by FlowJo software. Data were collected from three independent experiments and were analyzed by two-way ANOVA with Sidak’s post-tests. The bar graphs represent mean ± SEM *p-value <0.05, ****p-value <0.0001, n = 3.

### HMGB1 depletion modulates the DDR network following cisplatin treatment in human ovarian cancer cells

Cisplatin induces DNA intrastrand and interstrand crosslinks, resulting in cytotoxicity [10, 11]. These cisplatin-DNA lesions, especially ICLs, are highly cytotoxic, can stall DNA replication forks, and can result in the formation of DNA double-strand breaks (DSBs) [54, 55]. These cisplatin-induced DNA lesions can be recognized by the DDR machinery and then processed by several DNA repair proteins/pathways [13, 23, 24, 56]. To investigate how HMGB1 affects cisplatin sensitivity, we determined whether HMGB1 inhibition modulated DDR signaling and DNA repair capacity following cisplatin treatment in A2780 and CP70 cells. To do this, siHMGB1-treated and siNT-treated A2780 and CP70 cells were incubated with cisplatin at their IC50 concentrations for 24 h. Cell pellets were harvested at 0, 24, 48, and 72 hours post cisplatin treatment and subsequently subjected to western blot analysis to measure the levels of DDR proteins, including ATM, phospho-ATM (p-ATM), ATR, phospho-ATR (p-ATR), CHK1, phospho-CHK1 (p-CHK1), CHK2, phospho-CHK2 (p-CHK2) and several DNA repair proteins (**Figure 3A, B, Supplemental Figure 1A, B**). The densitometry Fiji software was used to semi-quantify the western blot images (**Figure 3C, D, Supplemental Figure 1C, D**). The band intensities were normalized to beta-actin as a loading control. Then, the normalized intensity values at each timepoint in the HMGB1-depleted and siNT groups were compared to the 0-hour value in the siNT group. The relative fold differences at each time point between the two groups were subsequently assessed to explore the impact of HMGB1 depletion on the levels of selected DDR and DNA repair proteins. Interestingly, HMGB1 depletion resulted in a reduction of total CHK1 levels in both A2780 and CP70 cells (NTC groups), and an increase of total CHK2 in CP70 cells, suggesting a role for HMGB1 in cell cycle regulation and/or a spontaneous background activation of DNA damage responses in cells. In addition, HMGB1 depletion in cisplatin-treated A2780 cells resulted in significant increases in the levels of p-ATM, p-ATR and p-CHK2 immediately after cisplatin treatment (0 hour), and reduced levels of p-CHK1 (at 0 and 24 hours). The total CHK1 level was also reduced at 0 hours and 24 hours. In cisplatin-resistant CP70 cells, HMGB1 depletion resulted in a similar, but later onset of alterations in response to cisplatin treatment, including increased levels of p-ATM (24 and 48 hours), p-ATR (24 hours) and p-CHK2 (24 hours), reduced CHK1 levels at all time points, and a reduction of the p-CHK1 level at 24 hours (**Figure 3C, D**). These results indicate that HMGB1 depletion enhanced the ATM/CHK2 pathway and attenuated the ATR/CHK1 pathway in response to cisplatin treatment, and the impact was more profound, but with a slower onset in the cisplatin-resistant CP70 cells compared to the A2780 cisplatin-sensitive cells.

**Figure 3.**
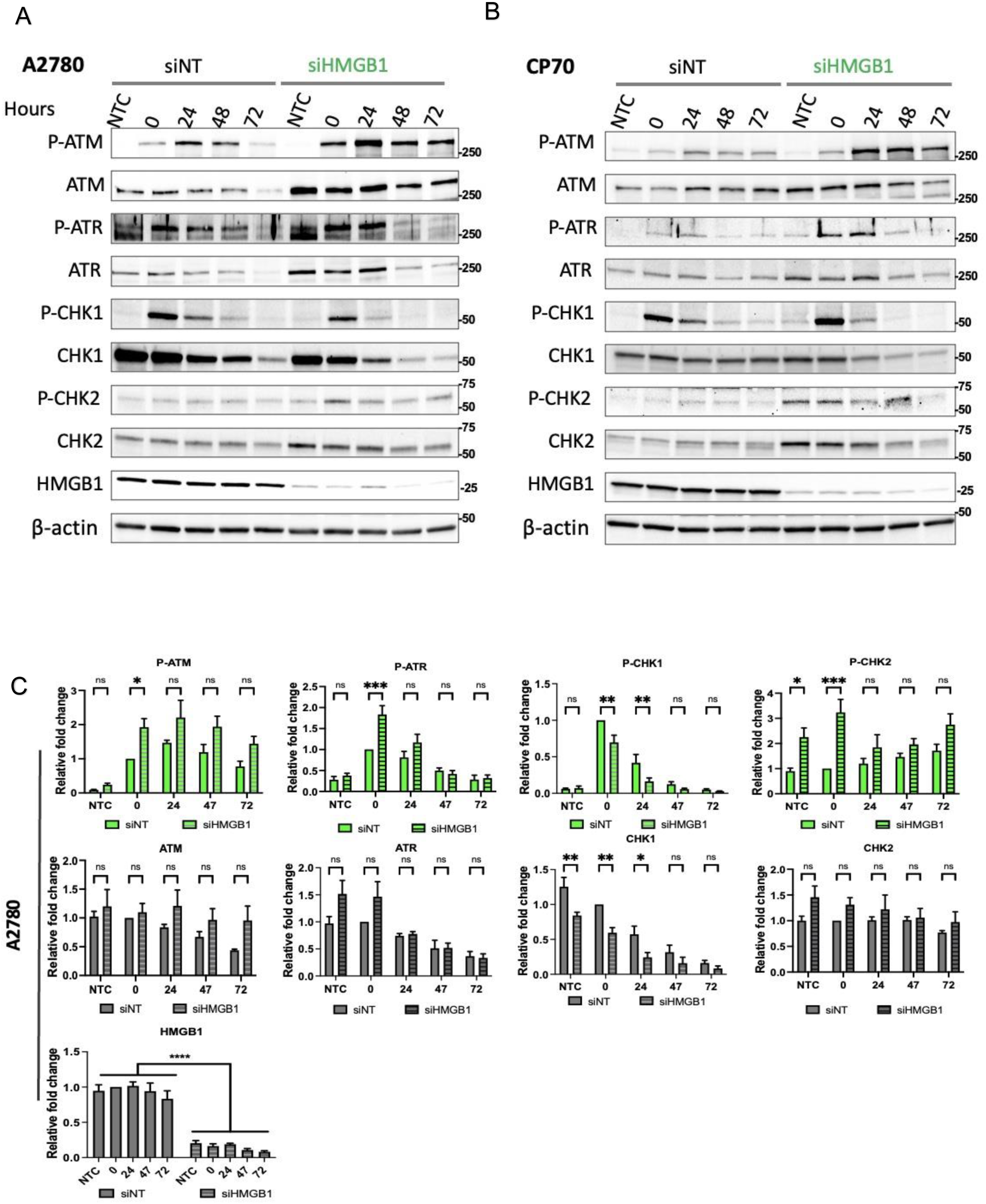
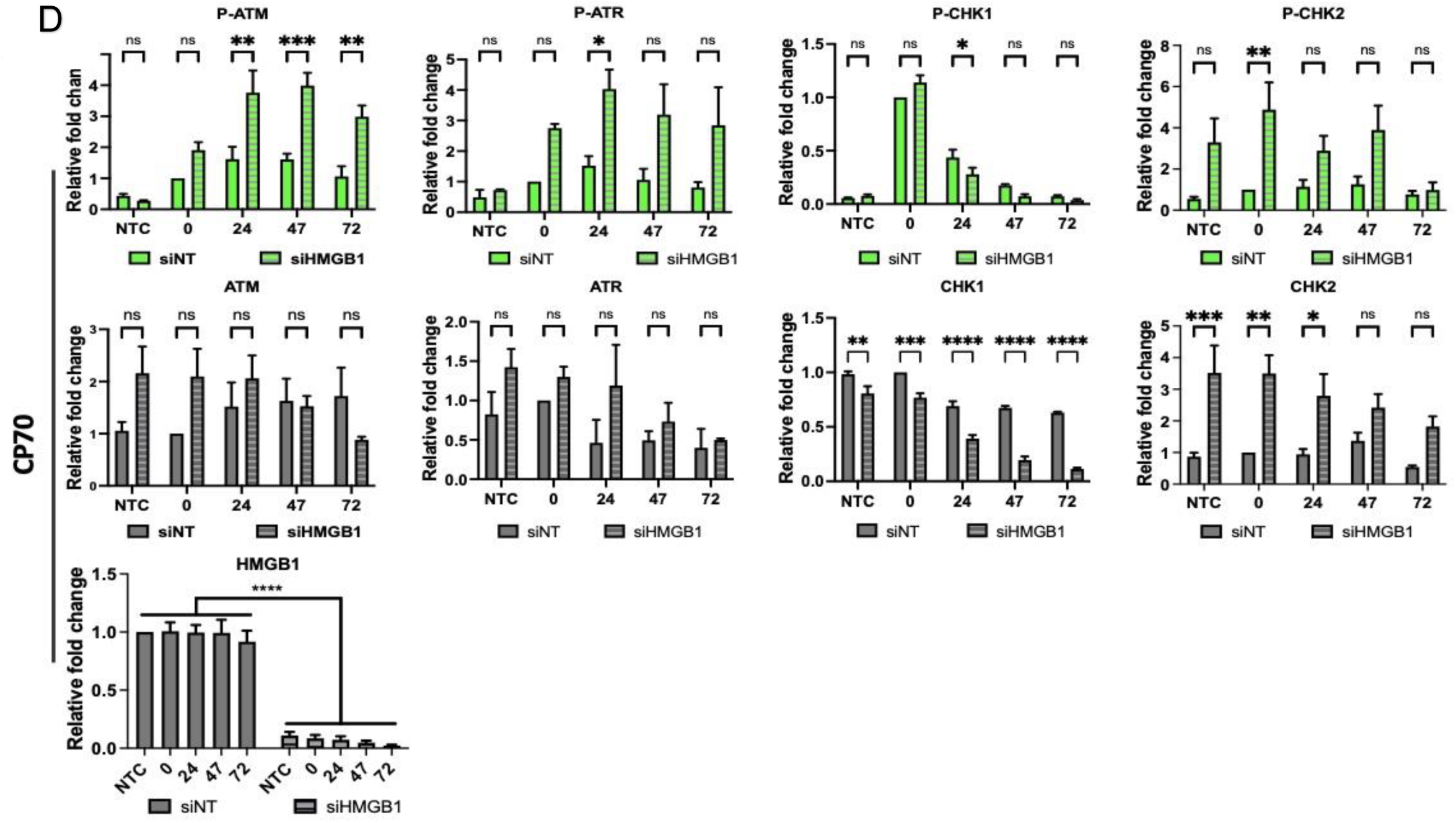
HMGB1 depletion enhances cisplatin sensitivity via altering DNA damage response proteins following cisplatin treatment in human ovarian cancer cells. Representative western blot panels showing the levels of DDR proteins in non-targeting siRNA (siNT) and HMGB1-targerting siRNA (siHMGB1) groups following cisplatin treatment in **A,** A2780 and **B,** CP70 cells. Cells were exposed to cisplatin-containing media at the IC50 concentration for 24 hours and allowed to recover in fresh media. Cell collection was performed at 0, 24, 48, and 72-hours post-cisplatin treatment, followed by protein extraction and immune blotting analysis. β-actin was used as the loading control. Semiquantitative analysis of the relative fold changes in DDR proteins are graphically depicted in **C,** A2780 and **D,** CP70 cells. Phosphorylated and total protein levels of ATM, ATR, CHK1, and CHK2 were determined over the time course at 0, 24, 47, and 72-hours post-cisplatin treatment for siNT and siHMGB1 groups in both cell lines. The band densitometric intensities were obtained using Image J software. Data were normalized to β-actin, and then to the timepoint 0 hour in the siNT group for each protein. Data are shown as mean ± SEM from at least three independent experiments. Statistical analysis was performed using original two-way ANOVA with Sidak corrections. *p-value <0.05, **p-value <0.01, ***p-value <0.001, ****p-value <0.0001, ns = not significant.

The ATM/CHK2 pathway is activated in response to DSBs and is critical in ICL processing [20, 57, 58]. Thus, we further examined the impact of HMGB1 depletion on the levels of the DSB marker, γ-H2AX and key proteins involved in downstream ICL repair, such as NER and DSB repair proteins [59–61] across different time points following cisplatin treatment (**Supplementary Figure 1**). As expected, γ-H2AX levels were significantly increased from 0-72 hours following cisplatin treatment in A2780 cells. However, HMGB1 depletion diminished this increase in γ-H2AX levels at the 48-hour and 72-hour time points following cisplatin treatment (**Supplementary Figure 1C**). In contrast, HMGB1 depleted CP70 cells exhibited elevated levels of γ-H2AX in the absence of cisplatin treatment, compared to the siNT cells, suggesting a role for HMGB1 in DSB formation and/or repair (**Supplementary Figure 1D**). Cisplatin treatment increased the γ-H2AX levels in the siNT groups, with no further significant increase in HMGB-depleted CP70 cells and no significant differences between siHMGB1 and siNT groups were observed at the later time points. Furthermore, the depletion of HMGB1 caused minimal impact on DNA repair protein levels in A2780 cells following cisplatin treatment. However, in the CP70 cells, we detected a significant induction in ERCC1 and NBS1 levels at NTC, 0 hours, and 24 hours post-treatment in HMGB1-depleted cells, which correlated with increased γ-H2AX levels. These results suggest a role for HMGB1 in modulating DNA repair capacity via regulating the levels of ERCC1 and NBS1 in response to cisplatin treatment. Overall, these findings suggest that HMGB1 depletion alters both replication stress and DSB repair pathways, further driving the cell toward apoptosis and enhancing cisplatin sensitivity.

### HMGB1 depletion facilitates cisplatin-induced DNA damage formation and reduces the efficiency of cisplatin-DNA adduct removal in ovarian cancer cells

Given that HMGB1 is a multifunctional protein, including its involvement in modulating DNA structure and participating in DNA repair processes [26, 31, 62], we aimed to determine whether HMGB1 depletion increases cisplatin sensitivity in CP70 cells via mediating the induction and/or repair of cisplatin-DNA adducts. To test this, the A2780 and CP70 cells were transfected with either HMGB1-targeted (siHMGB1) or non-targeting (NT) siRNA, followed by cisplatin treatment at half the IC50 concentrations for 24 hours. The cells were subsequently allowed to recover and harvested at 0, 24, and 96 hours post-treatment and genomic DNA was extracted and purified. A monoclonal antibody specific for unrepaired cisplatin-modified DNA [63, 64] was used in an immuno-slot blot assay to quantify the levels of cisplatin-DNA adducts in genomic DNA in the samples over time. Representative slot blot images are shown for the A2780 (**Figure 4A**) and CP70 cells (**Figure 4D**). The results revealed a significant increase in cisplatin-DNA adduct levels immediately at 0 hours and persisted at 24 hours post-cisplatin treatment in the HMGB1-depleted A2780 and CP70 cells compared to the control (**Figure 4B, E**). However, by 96 h post-treatment, the differences in adduct levels between the two groups were no longer significant (**Figure 4B, E**). The repair of the cisplatin-DNA adducts was measured as the percent of cisplatin adducts remaining over time relative to the amount of adducts identified at the 0-hour time point (**Figure 4C, F**). Although there were differences in the amount of initial damage, the repair of cisplatin adducts in the HMGB1-depleted A2780 cells was similar to the control (**Figure 4C**). However, HMGB1 depletion in the CP70 cells led to a delay in cisplatin-adduct removal, resulting in a significant accumulation of unrepaired adducts at 24 hours post-cisplatin treatment relative to the control (**Figure 4F**). The impact of HMGB1 depletion in cisplatin adduct removal at the later time points was diminished in both cell lines, indicating the involvement of alternative repair pathways in resolving cisplatin-induced DNA damage in the absence of HMGB1. Collectively, our data suggest that HMGB1 depletion increases cisplatin adduct formation and impairs repair efficiency in an early response to cisplatin treatment in CP70 cells, potentially contributing to the enhanced cisplatin sensitivity observed in the HMGB1-depleted chemoresistant cell line.

**Figure 4.**
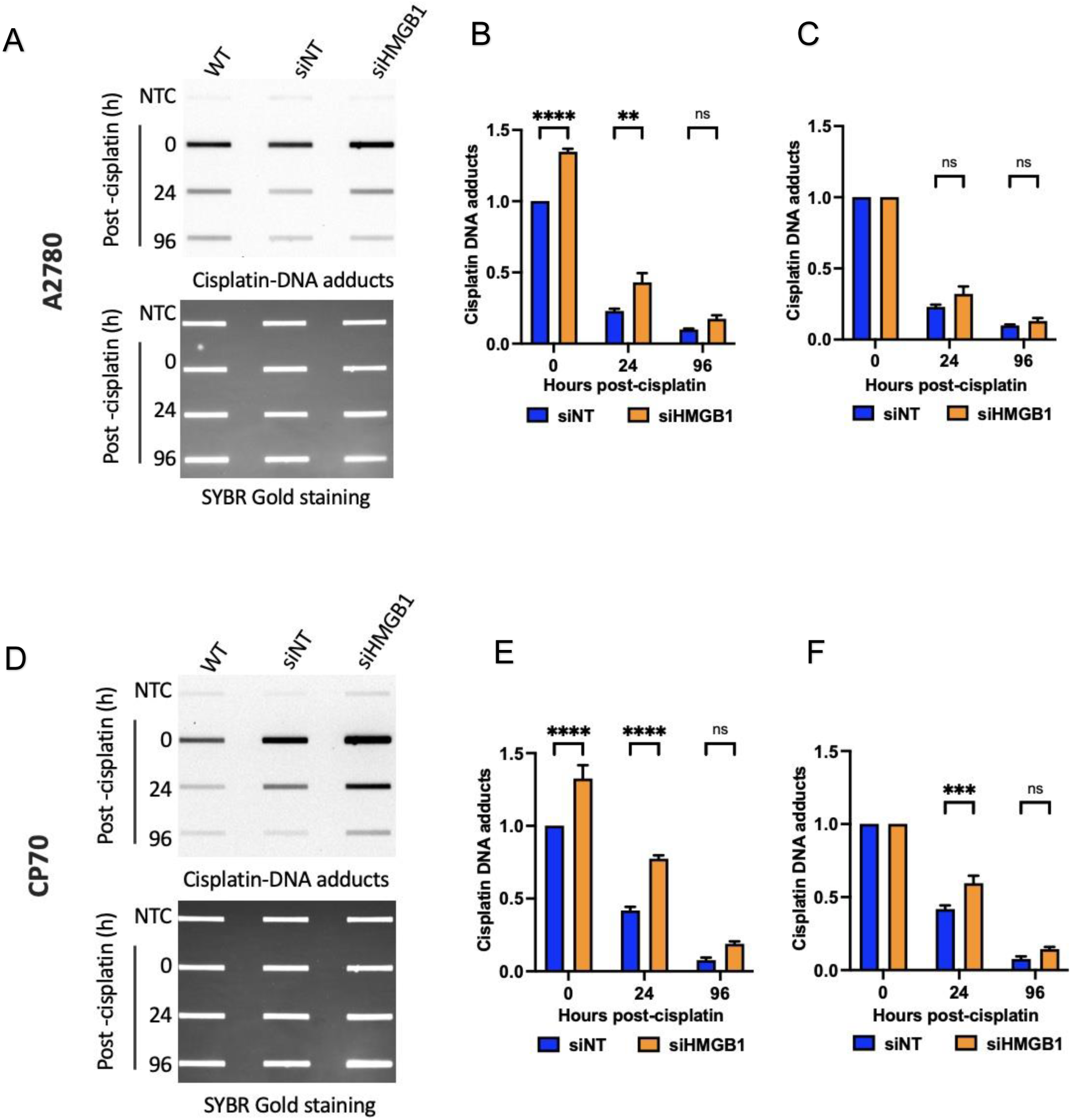
HMGB1 depletion modulates the formation and repair of cisplatin-induced DNA adducts in human ovarian cancer cells. Representative slot blot analysis to detect cisplatin-DNA adducts in **A,** A2780 and **D,** CP70 cells. Cells in the wildtype (WT), siNT and siHMGB1 groups were incubated with cisplatin media at 50% of the IC50 concentrations for 24 hours. The cells were recovered and harvested at 0, 24, and 96-hours post-cisplatin treatment. Genomic DNA was extracted and subjected to immunoblot analysis. Cisplatin-DNA adducts were detected using an anti-cisplatin modified DNA antibody. SYBR Gold staining was used to determine the amount of DNA loaded. An NTC (non-treated control) was included in each group to evaluate adduct formation in the absence of cisplatin. Quantifications of the slot blot data for cisplatin-DNA adduct formation are graphically represented in **B,** A2780 and **E,** CP70 cells. Band intensities were normalized to SYBR Gold intensities and then to the 0-hour timepoint in the siNT group. Quantification of cisplatin-DNA adduct removal is shown graphically in **C,** A2780 and **F,** CP70 cells. Band intensities at each timepoint in each group were normalized to the loading control, and then to the intensities at the 0-hour timepoint in that group to determine the timing of cisplatin adduct removal. Error bars represent mean ± SEM from at least three independent replicates. Statistical significance was determined by two-way ANOVA with Sidak’s multiple comparisons test. *p-value <0.05, **p-value <0.01, ***p-value <0.001, ****p-value <0.0001, ns = not significant.

### HMGB1 depletion stimulates cisplatin-DNA ICL induction and attenuates ICL repair efficiency in cisplatin-resistant ovarian cancer cancer cells

Cisplatin can interact with DNA to form intrastrand and interstrand crosslinks (ICLs) [10]. Both adducts contribute to the cytotoxic effects of cisplatin, with ICLs being the most toxic lesions [5]. Abnormalities in the processing of ICLs are known to modulate cisplatin sensitivity [60]. Thus, we aimed to investigate the association between cisplatin-induced ICL formation and removal using a modified alkaline comet assay [51], given the observed increase in cisplatin sensitivity in HMGB1-depleted ovarian cancer cells. The experimental approach for this analysis is outlined in **Figure 5A**. Samples from HMGB1-depleted (siHMGB1) and non-targeting siRNA-treated (siNT) cells were collected at 10, 24, 48, and 72 hours post-cisplatin treatment to assess ICL induction and repair over time. Cells were exposed to hydrogen peroxide (H_2_O_2_) immediately before lysis to induce a uniform amount of random single-strand breaks. ICL induction impedes the migration of fragmented DNA during electrophoresis, resulting in a decreased tail moment (DTM) compared to non-crosslinked H_2_O_2_-treated controls. The level of DTM correlates linearly with the frequency of cisplatin-induced ICLs in the genome [53]. Efficient ICL repair will result in the restoration of DTM over time. One hundred comets from each sample were analyzed using CometScore software to identify the mean Olive tail moment value, which was then used to calculate DTM and the percentages of ICL unhooking [53]. HMGB1 knockdown efficiency was assessed by western blotting (**Figure 5B**). Representative comet images for siHMGB1 and siNT-treated CP70 cells are shown in **Figure 5C**. Consistent with the published literature, we observed that cisplatin ICL induction was highest at 10 hours post-treatment in both the A2780 and CP70 cell lines. The data revealed that the level of DTM in HMGB1-depleted cells was significantly higher than the siNT cells at 10 and 72 hours post-cisplatin treatment, indicating that HMGB1 depletion facilitated cisplatin ICL formation and/or interfered with ICL removal, especially at the later time points (**Figure 5D**). Notably, we observed that HMGB1 depletion significantly increased ICL unhooking (∼48%, measured as the reduction of DTM at each time point from the maximum DTMs observed at 10-hour time point) compared to siNT (∼33%) (p <0.05) at 24 hours after cisplatin treatment (**Figure 5E**). The difference was no longer significant at 48 hours, and by the final time point at 72 hours, ∼73% of ICLs were unhooked in the siHMGB1 cells, significantly lower than that in the siNT cells (∼90%) (p <0.05) (**Figure 5E**), indicating that the removal rate of cisplatin ICLs was decreased in the absence of HMGB1. HMGB1 depletion did not result in significant differences in DTM and % ICL unhooking over a 48-hour time course in the A2780 cells (**Supplementary Figure 2A, B, C)**. These findings suggest that the depletion of HMGB1 modulates the formation and repair of cisplatin-induced ICLs, contributing to the increased cisplatin sensitivity observed in the CP70 cells.

**Figure 5.**
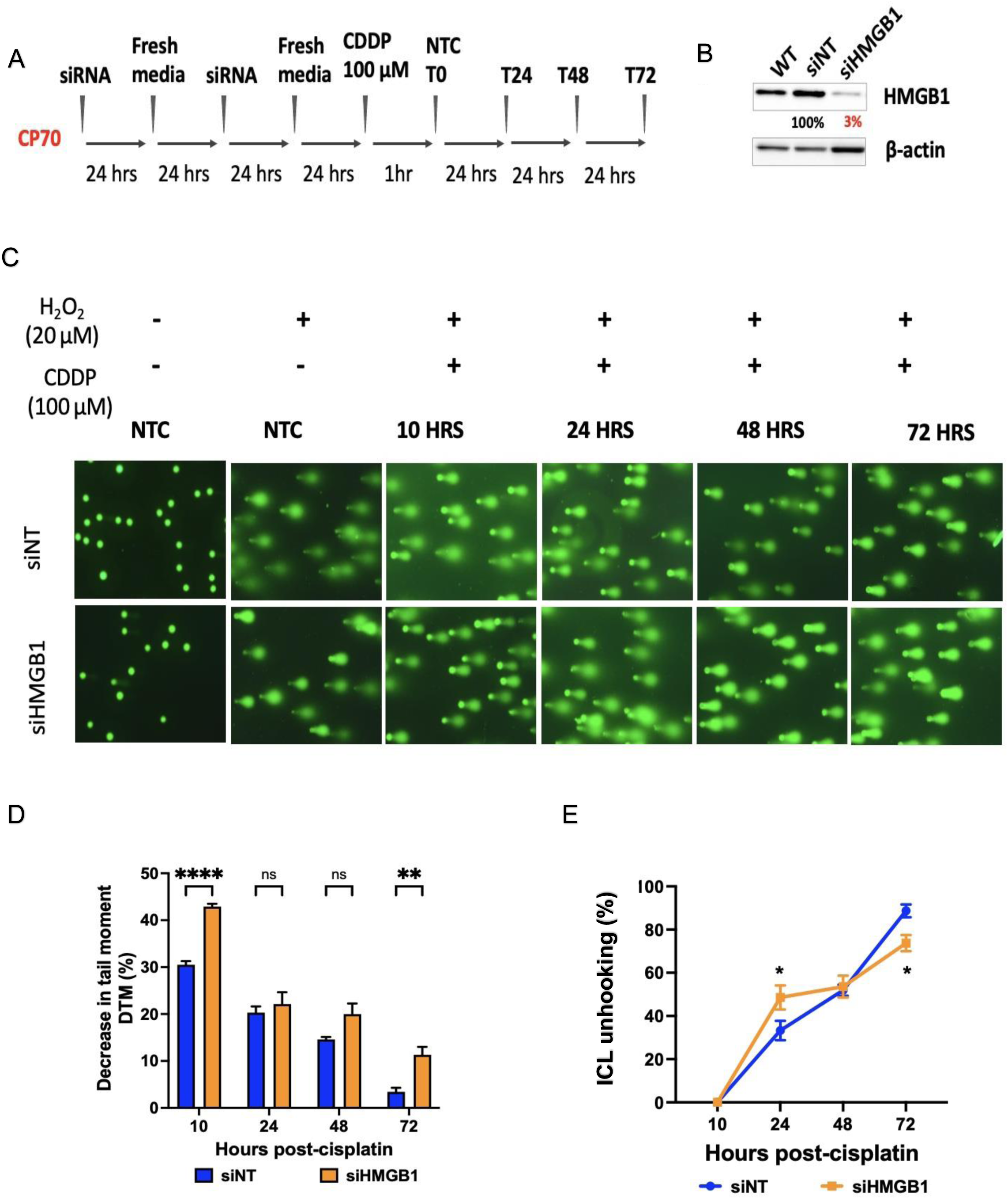
Modulation of cisplatin-induced ICL formation and repair post-cisplatin treatment in HMGB1-depleted chemoresistant ovarian cancer cells. A, Schematic overview of the experimental design to evaluate the impact of HMGB1 depletion on the processing of cisplatin - induced ICLs in CP70 cells using a modified alkaline comet assay (Refer to the Methods section for details). **B,** HMGB1 knockdown efficiency in CP70 cells was validated by western blot analysis. β-actin served as a loading control. **C,** Representative comet images acquired via CometScore 2.0 software showing the extent of DNA damage in siNT and siHMGB1 cells. **D, E,** Quantification of the modified alkaline comet assay results demonstrating ICL induction and repair in siNT and siHMGB1 cells. Panel D shows the decrease in the tail moment (DTM) at each time point after cisplatin treatment, where compared to the siNT cells, HMGB1-depleted cells showed significantly higher DTMs at 0 hour and 72-hours post-cisplatin treatment, reflecting the increase in adduct formation and/or the reduction in repair activity in the response to cisplatin treatment. Panel E shows the amount of cisplatin-ICL repair, graphed as the percentage of unhooking of ICLs over time relative to the 10-hour peak value in HMGB1-depleted and siNT CP70 cells. The data represent the mean ± SEM from three independent experiments. Statistical significance was calculated using original two-way ANOVA with Sidak’s post test. *p-value <0.05, **p-value <0.01, ***p-value <0.001, ****p-value <0.0001, ns = not significant.

## Discussion

Ovarian cancer is the ninth most common malignancy in the USA, with the highest mortality rate among gynecologic cancers [2, 65]. First-line chemotherapeutic treatment for ovarian cancer includes cisplatin. While cisplatin initially exhibits high antitumor efficacy, acquired resistance remains a significant challenge, limiting long-term treatment outcomes [4, 16]. Therefore, it is critical to understand the mechanisms underlying cisplatin resistance. The mechanisms of cisplatin resistance, while not fully understood, appear to be complex and involve multiple biological pathways [66]. Among these pathways, cisplatin resistance can be driven by upregulated DDR signaling and DNA damage repair activity (e.g. NER, MMR, and HR) [66–75]. These repair mechanisms allow cancer cells to efficiently remove cisplatin-induced DNA adducts, particularly ICLs, thereby escaping the cytotoxic effects of cisplatin. Additionally, resistance can also develop due to altered DNA methylation [76], dysregulated cell cycle control, impaired apoptosis [77], and activation of the epithelial-mesenchymal transition (EMT) pathway [78, 79]. In this study, we focus on the modulation of DDR signaling and DNA repair processes as a function of the HMGB1 protein in response to cisplatin treatment. HMGB1 is a non-histone DNA binding protein that interacts with cisplatin-modified DNA and plays important roles in chromatin remodeling and DNA repair processes [27, 37, 80]. Our study investigates the potential roles of HMGB1 in mediating cisplatin sensitivity and the underlying mechanisms.

Results from clonogenic survival and flow cytometry assays revealed that HMGB1 depletion resensitized chemo-resistant CP70 ovarian cancer cells to cisplatin (**Figure 1**) and promoted apoptosis upon cisplatin treatment (**Figure 2**). Furthermore, upon cisplatin exposure, DDR signaling and DNA repair activities were altered in HMGB1-depleted human ovarian cancer cells (**Figure 3, Supplementary Figure 1**). We further demonstrated that depletion of HMGB1 significantly facilitated the formation of cisplatin-DNA adducts but impaired adduct removal, especially in the chemoresistant ovarian cancer cells (**Figures 4** and **5**), consistent with the observed increase in cisplatin sensitivity.

Cisplatin can result in the formation of DNA intrastrand and interstrand crosslinks, which can lead to stalled replication forks and DSBs [54, 55]. The ATR/CHK1 pathway is critical in maintaining replication fork stability and cell cycle progression [18, 58]. The increase in p-ATM and p-ATR in HMGB1-depleted cells suggests that HMGB1 depletion increases unresolved cisplatin-induced DNA damage, triggering more robust cellular responses to replication stress and DSBs. Interestingly, while the ATR downstream effector kinase CHK1 was suppressed in HMGB1-depleted cells, the ATM effector kinase CHK2 was significantly activated, indicating that the absence of HMGB1 facilitated the CHK2-mediated response to cisplatin treatment, perhaps due to DSB formation and/or unknown regulatory mechanisms that selectively disrupt the ATR/CHK1 pathway. Consistently, while the depletion of HMGB1 did not impact total CHK2 levels in the A2780 cells, it was significantly increased in the CP70 cells following cisplatin treatment. Notably, the levels of ATM and ATR were not significantly different relative to the control in HMGB1-depleted cells. As the activation of the ATM/CHK2 pathway is implicated in facilitating DSB repair [20], we examined the levels of the DSB marker, γ-H2AX, following treatment with cisplatin. In the HMGB1-depleted A2780 cells, the levels of γ-H2AX were significantly reduced at 48 h and 72 h following cisplatin treatment, suggesting that HMGB1 depletion decreased cisplatin-induced DSBs and/or enhanced DNA damage repair activity in this cell line. In contrast, in the cisplatin-resistant CP70 cells, we found that cisplatin-induced γ-H2AX levels were significantly increased in the NTC and the 0-hour timepoints in HMGB1-depleted cells post-treatment. These results suggest that HMGB1 depletion exacerbated cisplatin-induced DNA damage in an early response to treatment in the CP70 cells, which may be related to the function of HMGB1 in modulating chromatin structure during DNA repair [26]. Taken together, our findings suggest that HMGB1 depletion altered DNA damage responses and repair activities in response to cisplatin treatment, consistent with the increase in cisplatin-induced cytotoxicity in the chemoresistant CP70 cells.

Given the established role of HMGB1 in chromatin remodeling and DNA repair, we further investigated whether HMGB1 depletion impacts the formation and processing of cisplatin-DNA adducts. Results from the slot blot assays (**Figure 4**) revealed that HMGB1-depleted CP70 cells accumulated significantly higher levels of cisplatin-DNA adducts at early time points post-treatment compared to the HMGB1-containing CP70 cells. The delay in adduct removal suggests that HMGB1 is involved in early DNA repair processes, particularly in chemoresistant cells. Previous results from our laboratory demonstrated that HMGB1 acts as an NER co-factor and interacts with RPA, XPA, and XPC in binding to ICLs [39, 41]. The persistence of adducts at 24 hours post cisplatin treatment in the current study suggests that HMGB1 impacts the efficient of recognition and/or processing of these lesions, consistent with its established role in NER.

The modified alkaline comet results revealed a role for HMGB1 in the repair of cisplatin ICLs, the most toxic form of cisplatin-induced DNA adducts. We observed a significant decrease in the tail moment (DTM) at 10 and 72-hours post-treatment in HMGB1-depleted CP70 cells, indicating a substantial increase in ICL formation and/or an impairment in ICL repair. Notably, while early ICL unhooking appeared to occur more rapidly in HMGB1-depleted cells at 24 hours, these cells failed to maintain efficient repair by 72 hours, suggesting a role for HMGB1 in facilitating efficient and/or sustained DNA damage repair.

Our findings from slot blot and modified alkaline comet assays in CP70 cells showed that HMGB1 depletion enhanced cisplatin-induced DNA damage formation and/or reduced the efficiency of cisplatin-DNA adduct removal, resulting in an accumulation of DNA lesions and decreased cell viability. These results align with our finding that the processing of UVC irradiation-induced DNA intrastrand crosslinks was significantly delayed in HMGB1 knockout mouse embryonic fibroblasts (MEFs) compared to wild-type MEFs [38]. Furthermore, HMGB1 knockout cells showed increased sensitivity to DNA-damaging agents, including UVC irradiation and psoralen-induced ICLs [38].

Previous studies have identified HMGB1 as a “repair-shielding protein,” binding to cisplatin-induced DNA adducts and protecting the damage from repair processes, thereby enhancing the cytotoxic effects of cisplatin. These effects have primarily been observed with cisplatin-DNA intrastrand crosslinks [81–83]. However, our published data and current findings suggest a different role for HMGB1, highlighting its function in facilitating the repair of both UVC- and cisplatin-induced intrastrand crosslinks [38]. Additionally, we have demonstrated that HMGB1 promotes the processing of psoralen- and cisplatin-induced ICLs across various cell types, modulating cell sensitivity to these DNA-damaging agents [38, 39, 41]. The discrepancy between our findings and the repair shielding hypothesis may be due to differences in experimental design, cell types, or the specific DNA repair pathways active in the cell lines, especially in cisplatin-resistant cells.

In conclusion, our data demonstrate that HMGB1 depletion sensitizes chemoresistant CP70 cells to cisplatin by impairing DDR signaling pathways and DNA repair processes, particularly the repair of cisplatin-DNA intrastrand and ICL adducts. These findings highlight the potential of targeting HMGB1 to overcome cisplatin resistance in ovarian cancer.

## Acknowledgments

We thank the Vasquez Lab for their support and valuable discussions over the years. This work was supported by NIH/NCI grants CA193124 and R01 CA093729 (to K.M.V.).

---

**Supplementary Figure 1.**
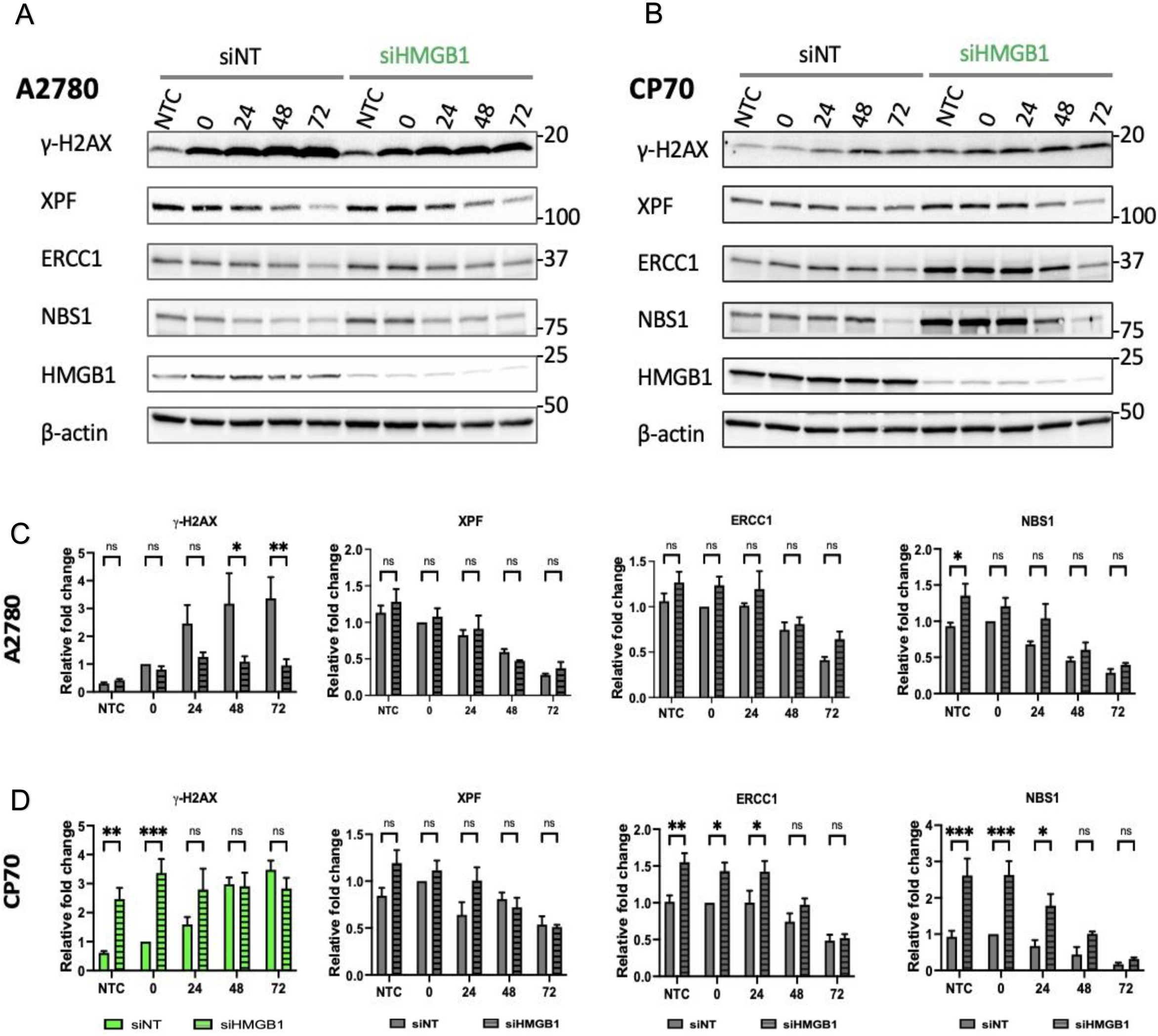
Impact of HMGB1 depletion on DNA repair protein levels in A2780 and CP70 cells in response to cisplatin treatment. Representative western blot panels for **A,** A2780 and **B,** CP70 cells. Cells were transfected with non-targeting siRNA (siNT) or HMGB1-targeted siRNA (siHMGB1) and then incubated with cisplatin for 24 hours before harvesting for western blot analysis. (See the Methods section for a detailed description), β-actin was used as a loading control. **C, D,** Quantification of normalized band intensities represents the relative fold changes in protein levels, normalized to the 0-hour timepoint in the siNT group. Error bars are mean ± SEM from at least three independent replicates. Statistical significance was analyzed using two-way ANOVA with Sidak’s multiple comparison test. *p-value <0.05, **p-value <0.01, ***p-value <0.001, ****p-value <0.0001, ns = not significant.

**Supplementary Figure 2.**
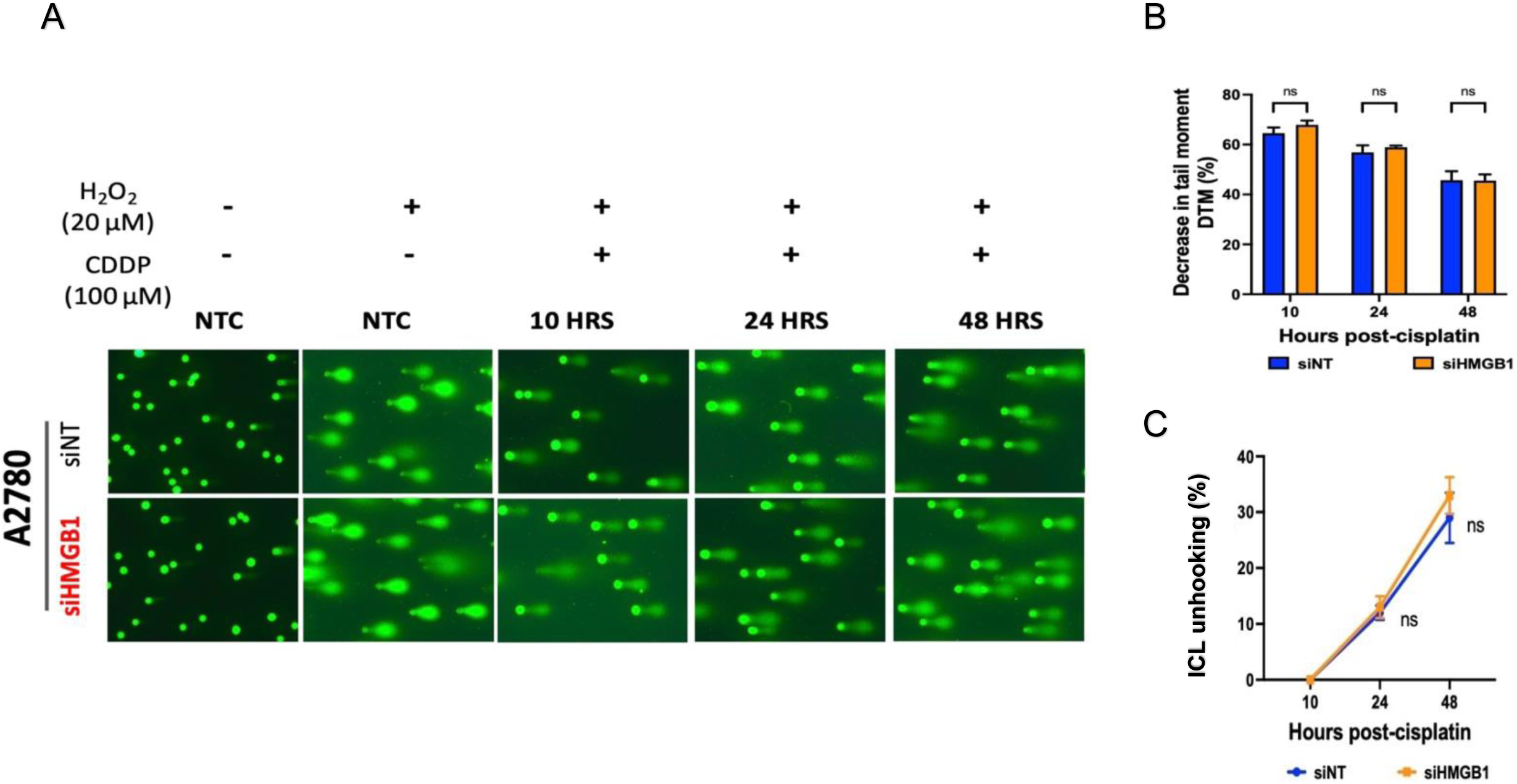
Cisplatin-ICL induction and removal were not affected by HMGB1 depletion in cisplatin-sensitive ovarian cancer cells. Similar alkaline comet experiments were performed in cisplatin-sensitive A2780 cells to evaluate the effects of HMGB1 depletion on cisplatin-induced ICL formation and processing in this cell line. The acquired images (**A**) and data analysis (**B**, **C**) showed non-significant differences in the formation of cisplatin-ICLs and the repair efficiency between experimental groups. Data represent the mean ± SEM of three independent experiments. Two-way ANOVA with Sidak’s post-test was used for statistical analysis. *p-value <0.05, **p-value <0.01, ***p-value <0.001, **p-value <0.0001, ns = not significant.

**Supplementary Table 1:**
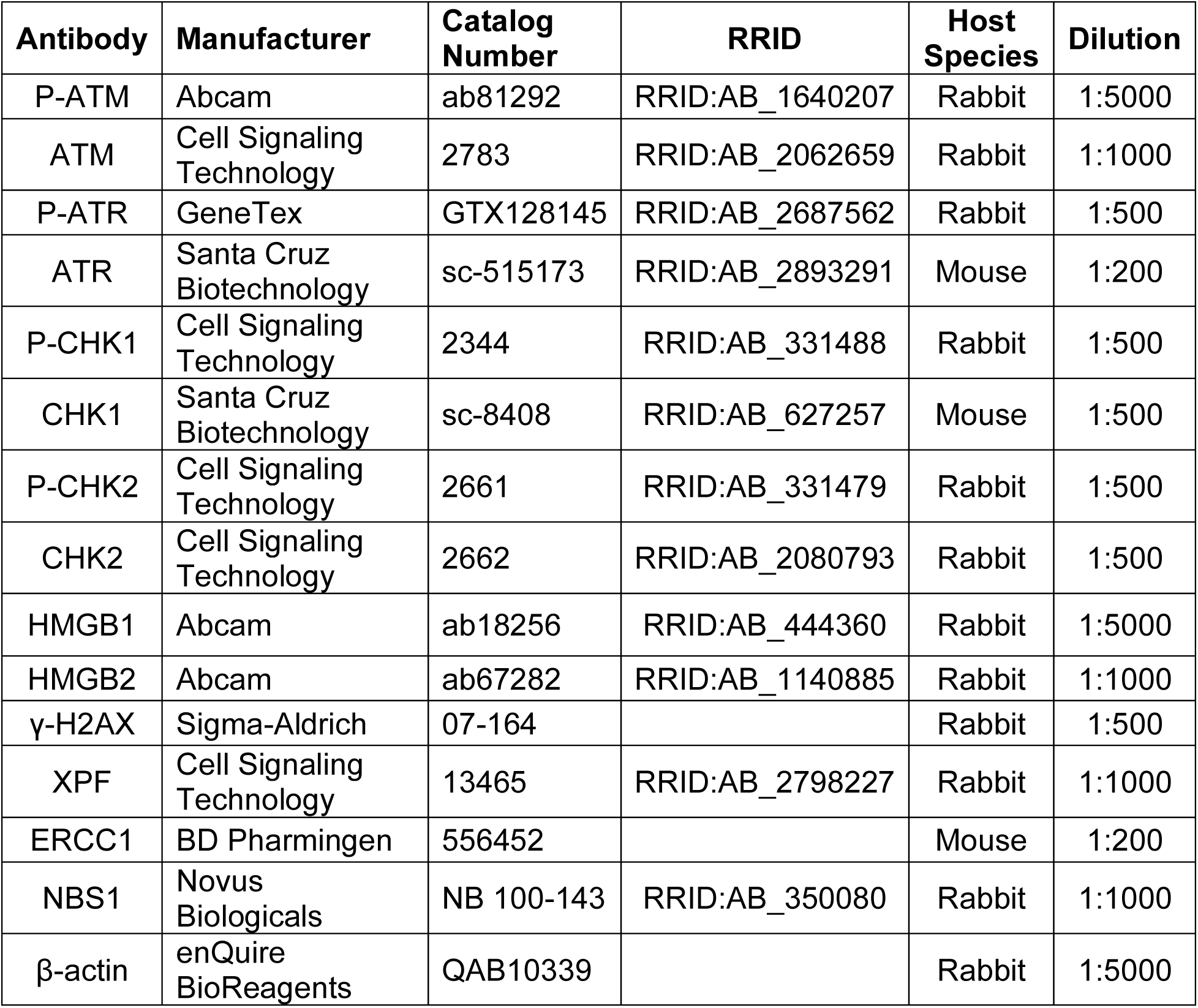
List of antibodies used in western blotting.

